# The assembly and efflux processes revealed by *in situ* structures of the ABC-type tripartite pump MacAB-TolC

**DOI:** 10.1101/2024.02.22.581608

**Authors:** Tong Huo, Wei Zheng, Zhili Yu, Yaoming Wu, Qiuyu Ren, Junli Cao, Zhao Wang, Xiaodong Shi

## Abstract

The MacAB-TolC tripartite efflux pump is widely distributed in Gram-negative bacteria, extruding antibiotics and virulence factors that lead to multidrug resistance and pathogenicity. This pump spans the cell envelope through its three components, an inner membrane ATP-binding cassette (ABC) transporter MacB, a periplasmic adaptor protein MacA, and an outer membrane protein TolC. However, the assembly process and efflux mechanism of the MacAB-TolC in cells remain unclear. Here, we resolve the *in situ* structures of *Escherichia coli* (*E. coli*) MacAB-TolC efflux pump by electron cryo-tomography and subtomogram averaging. In *E. coli* without antibiotic treatment, we observe a fully assembled MacAB-TolC pump with a weak constriction density in the middle of the MacA region. When *E. coli* cells were treated with erythromycin, we discovered the emergence of MacA-TolC subcomplexes, indicating flexible binding of MacB in the presence of an antibiotic substrate. This finding was further validated by *in vivo* crosslinking results. Together, our data present the *in situ* assembly process of the MacAB-TolC and derive a substrate-driven working model for ABC-type tripartite pump.

## Introduction

The multidrug resistance (MDR) of Gram-negative bacteria has become one of the foremost global public health threats. This resistance renders the inefficacy of commonly used antibiotics, posing an urgent challenge for clinical anti-infection treatment. Within Gram-negative bacteria, the tripartite efflux pumps spanning the cell envelope account for the active expelling of antibiotics and toxic compounds out of the bacterial cells. This process is a significant determinant of bacteria survival under antibiotic pressure, leading to the initiation of MDR^1^. MacAB-TolC represents one of the tripartite efflux systems, comprising the outer membrane protein TolC, the periplasmic adaptor protein MacA, and the inner membrane transporter MacB from the ATP-binding cassette (ABC) superfamily^2^. This pump actively extrudes various substrates, including macrolide antibiotics and virulence factors, thus contributing to both drug resistance and virulence phenotypes in *Escherichia coli* (*E. coli*) and other Gram-negative bacteria^3^.

The crystal structures of MacA^4^, TolC^5^, and MacB^6,7^ were all accessible through the Protein Data Bank (PDB). However, there are only cryo-electron microscopy (cryo-EM) single-particle structures of fusion-stabilized or disulfide bond-stabilized MacAB-TolC pump due to the difficulty in isolating the native full assembly through the purification procedure^8^. The failure to purify the fully assembled MacAB-TolC indicates low binding affinities among three components. Furthermore, it is essential to highlight that both the assembly and functioning processes of tripartite pumps require a cellular environment. All the above made capturing the intermediate states during the assembly and functioning of MacAB-TolC pump *in vitro* a huge challenge, leaving the assembly and functioning mechanism of this pump in living bacteria unclear.

TolC is the outer membrane component of several tripartite efflux pumps. In our previous study on another TolC-containing tripartite pump AcrAB-TolC, which belongs to the resistance-nodulation-cell division (RND) superfamily, we found that AcrA and AcrB first form a subcomplex and then recruit TolC^9^. This suggested a possible tripartite pump assembling path, which is periplasmic membrane fusion protein (AcrA) and inner member component of the pump (AcrB) could bind to each other first and then recruit the outer membrane component (TolC). However, whether all the TolC-containing tripartite pumps are assembled in the same sequence with TolC being recruited last is unclear. Besides, previous studies have shown that purified MacA alone can bind TolC^10,11^ as well as MacB^12^, and ATP binding could increase the affinity of the interaction between MacA and MacB^10^. Nevertheless, the impact of these interactions on the assembly of the tripartite pump and its functional state post-assembly remains unclear. Here, by employing electron cryo-tomography (cryo-ET) and subtomogram averaging, we resolve two different *in situ* structures of the MacAB-TolC efflux pump from *E. coli* cells in the native state and substrate transporting states. Our finding provides insights into the assembly and substrate-transport mechanisms of ABC-type tripartite efflux pump within its physiological cellular environment.

## Results

### Visualization of MacAB-TolC pumps in *E. coli* cell envelope

To visualize MacAB-TolC pumps in the *E. coli* cell envelope, we adopted the same strategy as our previous efflux pump studies to augment *in situ* MacAB-TolC concentration (Supplementary Figure 1)^9,13^. Next, we used cryo-ET to directly image the cells expressing *macA*, *macB,* and *tolC* genes, without any antibiotic treatment. In total, 70 tilt series were collected using a 300 kV microscope. The three-dimensional (3D) tomographic reconstruction showed that the bacterial envelope had numerous channel-like densities spanning through, indicating the presence of assembled MacAB-TolC (Figure 1). The visualization and following segmentation and labeling of the reconstructed densities present a uniform particle size, and the contrast of the image is consistent with our previous studies of AcrAB-TolC tripartite pump in bacteria^9,13^. Additionally, we also observed that the distance between the inner and outer membranes remained constant at the locations where the MacAB-TolC pumps occurred, implying that these assemblies may be constricting the periplasm (Figure 1A, B, D). In the tomograms, the densities of side-view particles appear as paired lines connecting the inner and outer membranes and spanning the entire cell envelope (Figure 1D). When viewed from the top, the particles appear as discernible circular features (Figure 1C).

**Figure 1.**
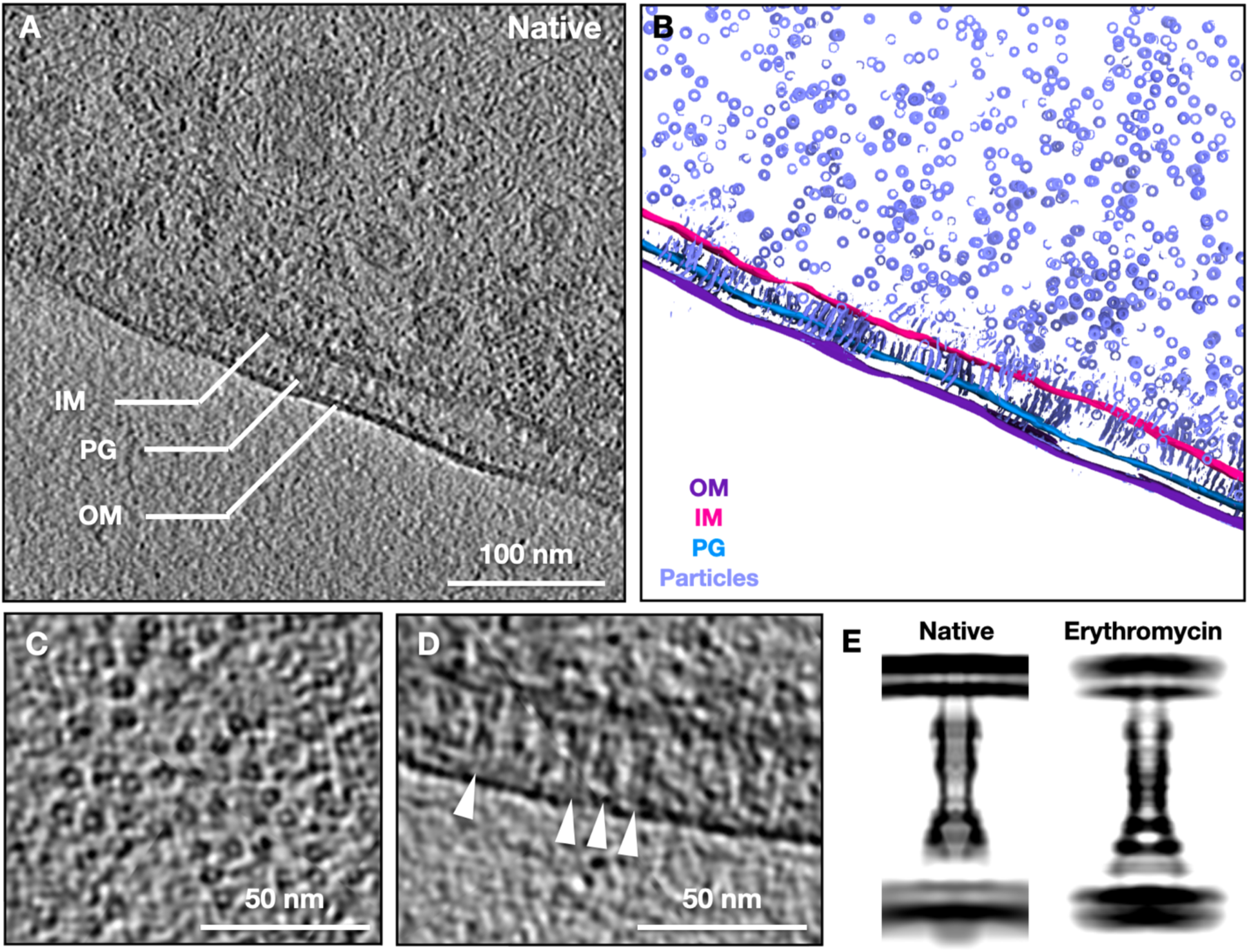
Visualization of the MacAB-TolC pump in the native *E. coli* cell envelope. A, a single Z slice from a tomogram of an *E. coli* cell expressing MacA, MacB, and TolC. B, a tomogram segmentation of A. C, zoom-in view of the MacAB-TolC top view particles. D, zoom-in view of the MacAB-TolC side view particles (indicated by white arrowheads). E, the 2D projection of the rotationally averaged density map of the *in situ* structure of MacAB-TolC in the native (left) and erythromycin-treated (right) *E. coli* cells. IM, inner membrane; OM, outer membrane; PG, peptidoglycan.

### *In situ* structure of the fully assembled MacAB–TolC complex

For subtomogram averaging, 5,183 particles were used, and the final refinement achieved 14 Å resolution (Supplementary Figure 2). The MacAB-TolC structure (PDB: 5NIK) was docked into the map, with the overall shape of MacB, MacA, and TolC structures well-fitted with the map. The density for the outer and inner membrane observed in this study is located at the TolC β-barrel domain and the MacB transmembrane domain, respectively, indicating that the MacAB-TolC pump spans the full periplasm (Figure 2A). The interior of the density corresponding to TolC viewed in a cross-section through the averaged map did not have any constriction site at the interface between MacA and TolC, which made TolC a fully open tunnel to the top of the outer membrane (Figure 2A). In contrast, we observed within the density corresponding to MacA a weak constriction sealing the pump channel, the position of which corresponds to the gating ring of MacA (Figure 2A)^8^. This indicates that MacA resembles a closed state under this experimental condition. The periplasmic domain (PLD) of MacB could be identified, which forms interactions with MacA. The nuclear binding domain (NBD) of MacB could also be visualized, indicating the presence of the MacB cytoplasmic part (Figure 2A).

**Figure 2.**
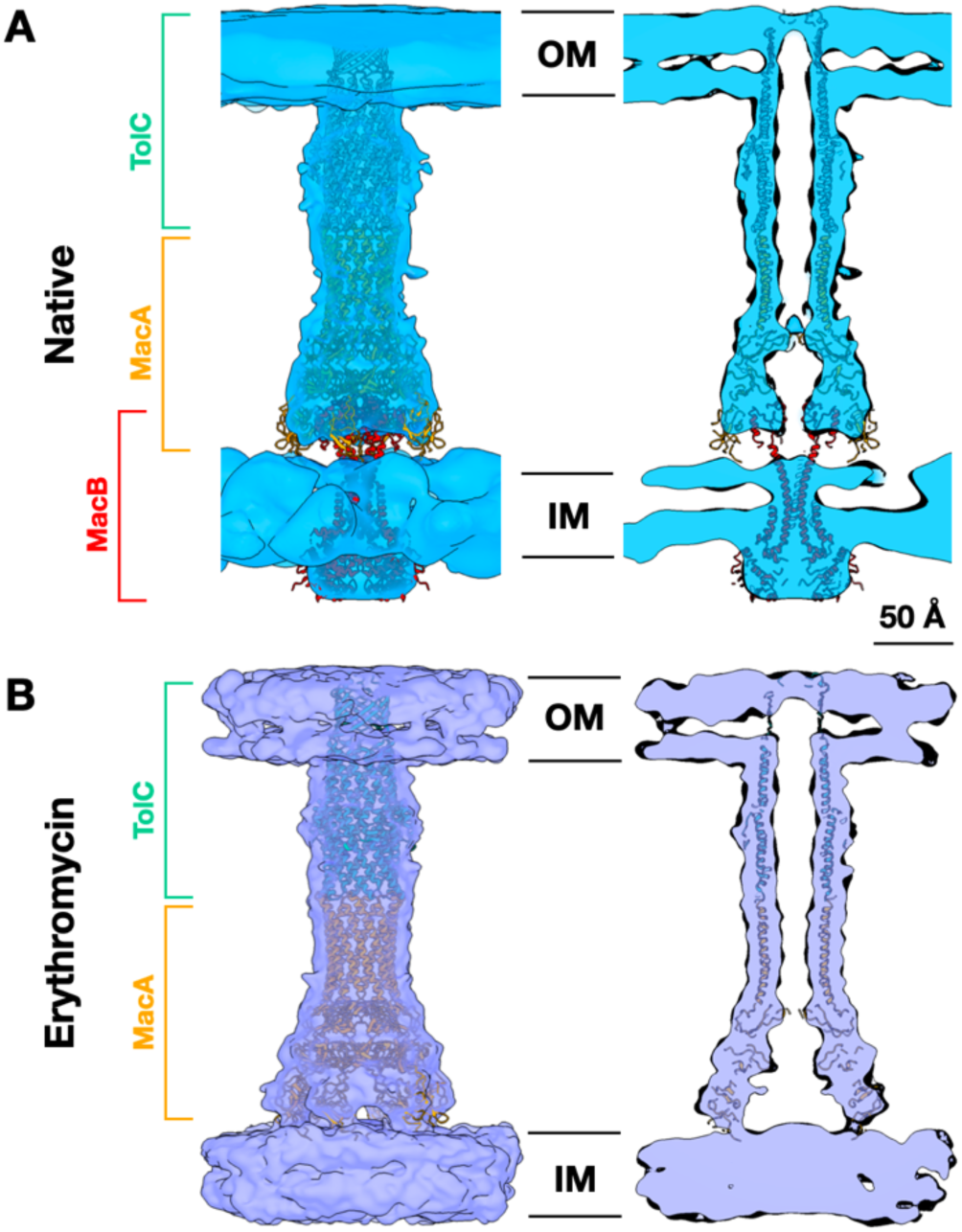
*In situ* cryo-ET structures of the assembled MacAB-TolC complex and MacA-TolC subcomplex. A, density maps of MacAB-TolC in the native state fitted with high-resolution MacAB-TolC cryo-EM model (PDB: 5NIK), and its central cross-section view. B, MacA-TolC in the adding erythromycin state fitted with the high-resolution MacAB-TolC (MacB-omitted) cryo-EM model (PDB: 5NIK), and its central cross-section view. IM, inner membrane; OM, outer membrane.

### Erythromycin facilitates the detaching of MacB from MacAB-TolC

To understand pump structural response under antibiotic pressure, we treated the MacAB-TolC-overexpressing *E. coli* cells with erythromycin and followed the same imaging pipeline as mentioned above. Erythromycin is the main substrate of MacAB-TolC and a representative of the macrolide antibiotics, which is one of the commonly used antibiotics in clinical treatment. Except for the treatment by erythromycin, all the other experimental processes are the same as those for native *E. coli* cells. The minimum inhibitory concentration (MIC) results indicate that the MacAB-TolC pumps overexpressed by two plasmids can transport erythromycin (Supplementary Table 1). The feature of 3D tomographic reconstruction resembles that of MacAB-TolC-overexpressing *E. coli* cells without erythromycin treatment (Supplementary Figure 3). However, the initial refinement shows that the density at the MacB region is weak compared to MacA and TolC in the complex, suggesting an occupancy variation in the dataset. Next, using a focused classification strategy in the subtomogram averaging process, we classified the entire dataset into two datasets and reconstructed two different conformations of the complex. One of the classes is consistent with our native structure, while the second class shows an absence in the MacB region, as the PLD domain is invisible (Figure 1E). We resolved the structure of the MacA-TolC subcomplex at around 12 Å (Supplementary Figure 2). The structure of MacA-TolC is consistent with the counterpart of that in the MacAB-TolC pump. Missing MacB resulted in the formation of an empty chamber between MacA membrane-proximal (MP) domains and the inner membrane (Figure 2B). These results suggest that erythromycin, as a substrate of MacAB-TolC, facilitates MacB to detach from the pump complex in cells.

Based on this result, we further validated whether the MacA-TolC could form a stable complex on the cell envelope without MacB. We overexpressed MacA and TolC but not MacB in *E. coli* cells and prepared the cryo-ET specimen using the same strategy. We collected 67 tilt series and performed tomogram reconstruction. Within the tomograms, we could observe pump-like features across the cell envelope, indicating the existence of the MacA-TolC subcomplex (Supplementary Figure 4). This observation confirmed that MacA and TolC can bind to each other without MacB and form a complex on the cell envelope, which indicates that the MacA-TolC subcomplex may be formed first, followed by the binding of MacB to assemble the full tripartite pump. Based on the MacA-TolC subcomplex we have obtained under the situation of erythromycin treatment and MacA-TolC overexpression, we propose that the association/disassociation of MacB to the MacA-TolC subcomplex triggered the switch of the activation status of MacAB-TolC.

### *In vivo* crosslinking results demonstrate the detaching of MacB from MacAB-TolC upon erythromycin treatment

To further investigate whether erythromycin facilitates the detaching of MacB from MacAB-TolC, we first performed a formaldehyde crosslinking experiment on MacAB-TolC-overexpressing *E. coli* cells. The result shows that upon erythromycin treatment, the amount of MacA-MacB-TolC crosslinked product was reduced (Supplementary Figure 5), implying that erythromycin could cause the dissociation of MacB and MacA-TolC. Next, we performed *in vivo* photo-crosslinking mediated by *p*-benzoyl-L-phenylalanine (Bpa) to determine the interactions between MacA and MacB within *E. coli* cells. Based on the single-particle cryo-EM structure of MacAB-TolC (PDB: 5NIK), a total of 10 key residues involved in the MacA-MacB interactions were chosen for Bpa substitution, which are N428, P430, M464, G465, Q466, and L495 on MacB and T53, V269, D271, and R343 on MacA. The immunoblotting result shows that only the Bpa substitution of D271 on MacA seriously affected the expression of the *macAB* operon (Supplementary Figure 6). When the *E. coli* cells expressing Bpa variants were subjected to UV radiation, we observed the cross-linked products of MacB^P430Bpa^, MacB^L495Bpa^, MacA^T53Bpa^, MacA^V269Bpa^, and MacA^R343Bpa^ (Supplementary Figure 7), suggesting that these five residues are crucial to mediate the interaction between MacA and MacB. Through MIC assay, we confirmed that the function of the above five Bpa variants was barely affected by Bpa substitution (Supplementary Table 2). According to the positions of Bpa substitutions in MacA and MacB, MacB^L495Bpa^ and MacA^V269Bpa^ were chosen as the final representatives to investigate the interaction between MacA and MacB (Figure 3A). After treatment with erythromycin, the crosslinked product of both MacB^L495Bpa^ and MacA^V269Bpa^, namely MacA-MacB^L495Bpa^ and MacA^V269Bpa^-MacB, decreased significantly (Figure 3B-3E). Taken together, our *in vivo* protein photo-crosslinking data indicate that the interaction between MacA and MacB was reduced upon erythromycin treatment, corresponding to the structural result that MacB detaches from MacA-TolC in the presence of erythromycin.

**Figure 3.**
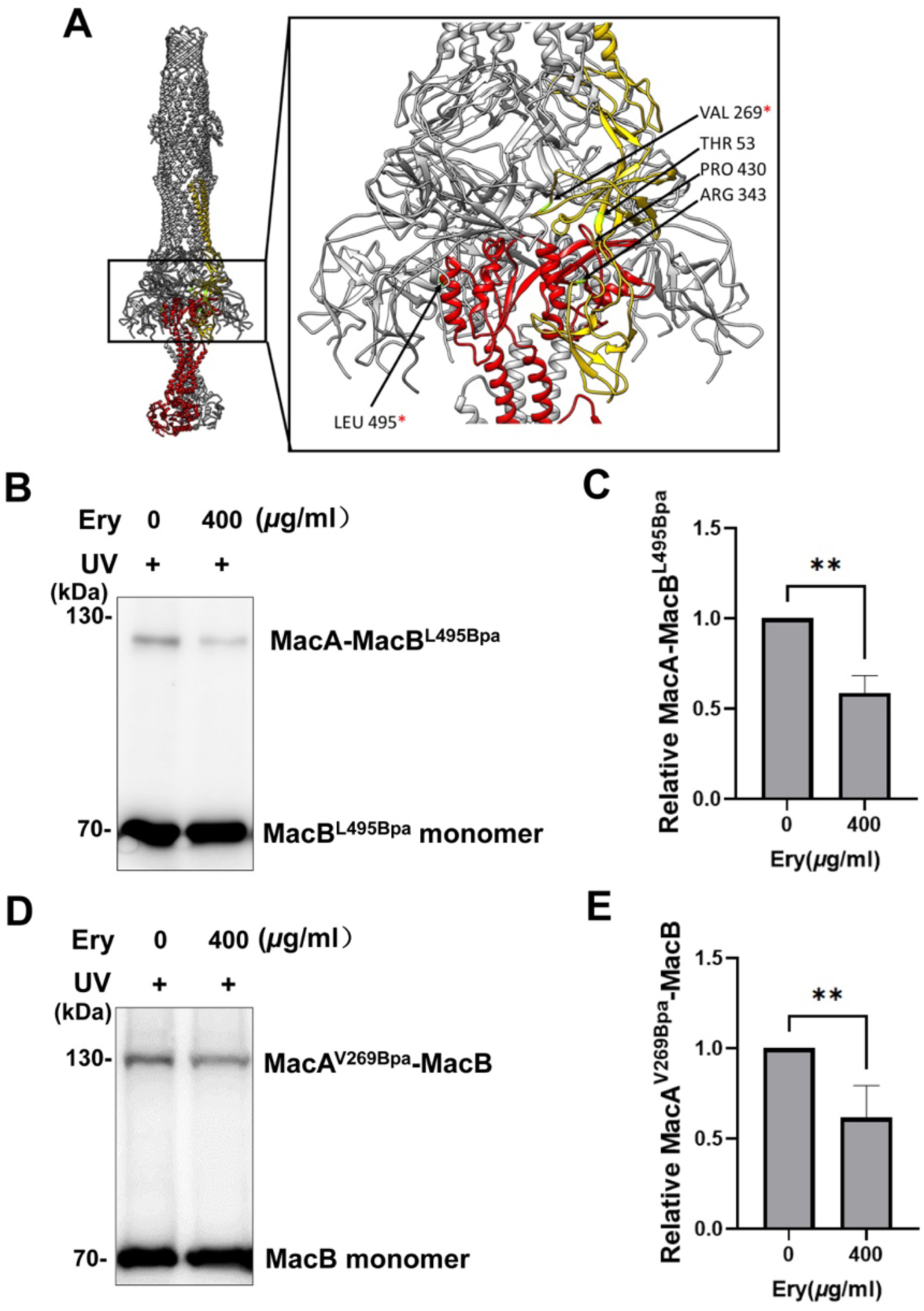
Immunoblotting results of the *in vivo* photo-crosslinked products of MacA/MacB Bpa variants upon erythromycin treatment. A, the positions of introduced Bpa in MacA/MacB are shown by red arrows. B and C, the *in vivo* photo-crosslinking results of MacB^L495Bpa^ after treatment with erythromycin (n = 3). D and E, the *in vivo* photo-crosslinking results of the MacA^V269Bpa^ after treatment with erythromycin (n = 4). Data are shown as the mean values ± SD. \*\**P* < 0.01. Ery, erythromycin.

## Discussion

In this study, we show the *in situ* structures of the MacAB-TolC efflux pump in *E. coli*, both in native and erythromycin-present states. Combining imaging results, protein crosslinking data, as well as previous works^10,12,14,15^, we propose a model for the *in situ* assembly and working mechanism of MacAB-TolC (Figure 4). Without substrates (apo state), MacA and TolC are highly likely to associate to form a stable bipartite complex in the cell. Next, ATP-bound MacB is recruited to form a stabilized tripartite MacAB-TolC pump. When the substrates are present, the binding of substrates induces a conformational change of MacB within the tripartite complex. In consequence, this alteration leads to a reduced affinity between MacA and MacB, resulting in the dissociation of MacB from the full pump. It has been suggested that ATP plays a crucial role in the assembly process of the MacAB-TolC efflux pump^10^, therefore, we speculate that the binding and hydrolysis of ATP could be an internal factor triggering the association and dissociation of MacB with the MacA-TolC subcomplex. Our finding that MacA-TolC subcomplexes exist as a stable entity in cells confirmed the previously proposed binding scheme from *in vitro* experiments^12,14,16^. Although previous biochemistry studies suggested MacA-MacB binding in the absence of TolC^10,12^, we did not observe any MacA-MacB bipartite subcomplexes stabilized in cells.

**Figure 4.**
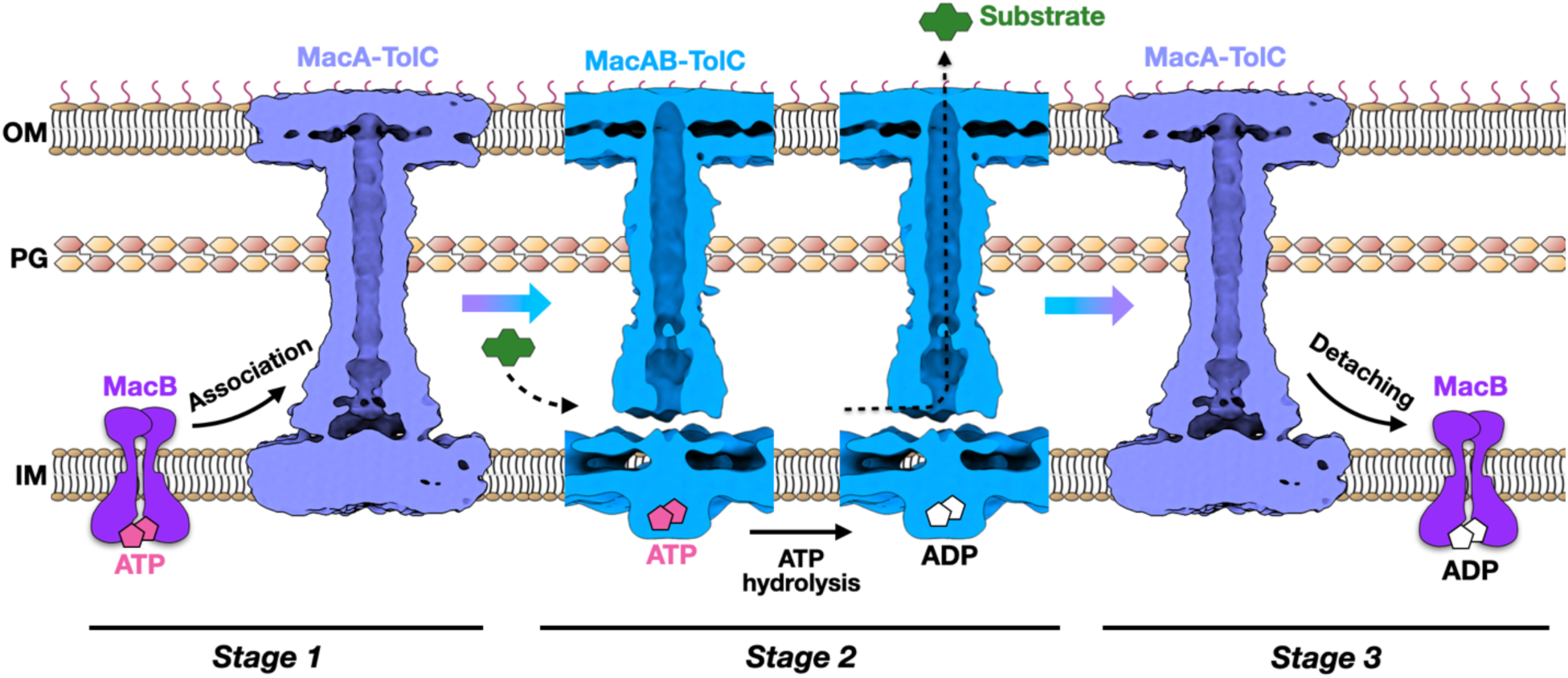
The assembly and working model of the ABC-type tripartite pump MacAB-TolC in *E. coli* cells. In the absence of substrates, MacA binds TolC to form the MacA-TolC subcomplex and then recruits ATP-bound MacB to assemble into the MacAB-TolC efflux pump. In the presence of substrates, MacAB-TolC hydrolyzes ATP to efflux substrates, and MacB dissociates from MacA-TolC subsequently. Note that the conformation of the gating ring in MacA in stage 1 is still unknown.

Overall, our results suggest that the assembly sequence of MacAB-TolC is distinct from AcrAB-TolC, in which the AcrA-AcrB bipartite subcomplex forms first and TolC is recruited at the appropriate time^9^. It is conceivable that the assembly sequence of different tripartite pumps may not necessarily follow a similar manner, although TolC is required for the above two pumps to be functioning. One of the possible reasons that causes MacAB-TolC and AcrAB-TolC to present different assembly sequences in the bacteria may be associated with their different energized mechanism between ABC and RND transporters in the pumps.

Our *in situ* data show that the fully assembled MacAB-TolC efflux pump on native *E. coli* membranes has a thin density constriction in the middle of the channel, the position of which corresponds to the gate ring in MacA. After treating pump-overexpressing cells with erythromycin, we captured the MacA-TolC subcomplex with the gate ring density in MacA disappearing or opening, and we infer that the gate is possibly in an open state, since the MacAB-TolC pump is actively pumping the erythromycin molecules in this state (Supplementary Table 1). These results indicate that the conformation of MacA in the tripartite MacAB-TolC complex differs from that in the bipartite MacA-TolC subcomplex, which is consistent with the *in vivo* proteolysis results that association with TolC and MacB changed the conformation of MacA, with TolC sensitizing MacA to proteolysis and MacB protecting MacA from proteases^17^. In contrast, we find that TolC stays open in both the MacAB-TolC complex and MacA-TolC subcomplex. The electron microscopic study of the complex of MacA with the tip region of TolC α-barrel revealed that MacA induced opening of the TolC channel^11^. Taken together, these results suggest that the opening of the MacAB-TolC channel appears to be controlled by MacA rather than TolC, unlike the AcrAB-TolC efflux pump. Besides, it has been found that the binding of MacA to MacB stabilized the ATP-bound conformation of MacB *in vitro*^17^. Previous studies on MacB have shown that MacB could have two conformations with ATP-bound or ADP-bound^6,7^. We are unable to determine whether MacB in our *in situ* structure is ATP-binding due to resolution limitations. However, we speculate that it is likely to be in an ATP binding state, as *in vitro* experiments showed that ATP binding increased the affinity of MacA-MacB complex, while ADP had a negative effect on kinetics of MacAB interaction^10^, which is likely to be amplified in the real cell membrane environment.

These results provide new insights into the working mechanism for ABC-type tripartite multidrug efflux pumps and present this distinct assembly process as a reference for designing novel efflux pump inhibitors. Confirming the MacA-TolC bipartite complex as a primary intermediate in the assembly process presents a novel target for therapeutic design. Given its unique occurrence in bacteria, targeting this crucial intermediate state during assembly can yield lower side effects in therapeutic interventions. This insight opens avenues for developing more precise and effective strategies in combating bacterial resistance.

## Materials and Methods

### Bacterial strains, plasmids, and protein expression

*E. coli* BW25113-Δ*acrB* strain was kindly provided by Dr. Xinmiao Fu at Fujian Normal University (Fuzhou, China). *E. coli* BL21(DE3)-Δ*acrAB-tolC* strain was constructed by Ubigene Biosciences Co., Ltd (Guangzhou, China). The DNA sequence encoding *macAB* operon (*E. coli* K12 strain) was synthesized by General Biosystems Co., Ltd (Anhui, China) and cloned into pBAD with a hexahistidine tag at the C terminus of MacB, yielding pBAD-*macAB*-his. The DNA sequence encoding *macA* was synthesized by General Biosystems Co., Ltd (Anhui, China) and cloned into pETDuet, yielding pETDuet-*macA*. The pRSF-*tolC* plasmid was obtained from our previous study^9^.

For cryo-ET imaging experiments, to overexpress the MacAB-TolC efflux pump, *E. coli* BL21(DE3) cells were co-transformed with plasmids pBAD-*macAB*-his (ampicillin resistant) and pRSF-*tolC* (kanamycin resistant). Cells were cultured at 37°C in LB medium supplemented with 100 *µ*g/ml ampicillin and 50 *µ*g/ml kanamycin. When OD_600_ reaches 0.8, 0.2% L-arabinose was added to induce the expression of MacAB, at 20°C, 210 rpm for 24 h. Then TolC was induced by adding 0.5 mM isopropyl 1-thio-β-D-galactopyranoside (IPTG) at 20°C, 210 rpm overnight. To overexpress MacA and TolC, *E. coli* BL21(DE3) cells were co-transformed with plasmids pETDuet-*macA* and pRSF-*tolC*. Cells were cultured at 37°C in LB medium with 100 *µ*g/ml ampicillin and 50 *µ*g/ml kanamycin until an OD_600_ of 0.8 was reached, and protein expression was induced by adding 0.5 mM IPTG at 20°C overnight.

### Cryo-ET sample preparation

After induction, *E. coli* cells were harvested by centrifugation at 20,000 g for 8 min, washed once with PBS buffer, and then centrifuged again at 20,000 g for 8 min. Cells were then resuspended to an OD_600_ of 10. For the native state, cells are ready for freezing in this step. For the erythromycin treatment state, cells were then added with 400 *μ*g/ml erythromycin, and incubated at 37°C, 210 rpm for 30 min. For freezing, cells were mixed with 6 nm BSA fiducial gold (Aurion). The volume ratio of BSA fiducial gold to sample is 1:3. A 3 *μ*l droplet of the sample was applied to the freshly glow-discharged, continuous carbon film-covered grids (Quantifoil Cu R3.5/1, 200 mesh with 2 nm continuous carbon film) and plunged frozen using a Vitrobot Mark IV (FEI). Grids were stored in liquid nitrogen until required for data collection.

### Cryo-ET data collection and 3D reconstruction

The frozen-hydrated samples were imaged on a Titan Krios microscope with a Gatan K2 Summit direct electron detector camera. The magnification was 81,000 x, with a pixel size of 1.76 Å. All tilt series were collected using a 3-degree angular step, from −51 to +51 degrees, with a starting angle of −30 degrees. The defocus ranges from −1.5 *μ*m to −5 *μ*m, and the total dose for each tilt series is about 100 e^−^/Å^2^.

### Cryo-ET data processing

The raw frames were aligned using MotionCorr2^18^. For the native state MacAB-TolC dataset, 8,734 particles were manually picked from 36 good tomograms. After generating the initial model from scratch, 3D refinement was carried out using EMAN2^19^. The particle orientations from this refinement were visually examined using the evaluate refinement function in EMAN2, which maps the particles together with their orientations back to each tomogram, allowing an overview of the refinement quality. During this process, we observed that in some tomograms, most of the top view particles are incorrectly aligned, hinting at an insufficient signal-to-noise ratio in these tomograms. Particles from these tomograms were excluded from the following data processing, leaving 5,183 particles that were used for refinement from scratch again. Geometric features of the cell membrane were used to correct particle orientations^20^, followed by one round of local refinement. The C3 symmetry was applied to all the refinements. A 14 Å density map was obtained, which could be fitted by the structure (PDB: 5NIK).

For the dataset of MacAB-TolC under treatment of erythromycin, a similar process was conducted. 877 particles were picked manually from 68 good tomograms and extracted, which were used for generating the initial model from scratch. After the initial model was built, one round of refinement using 2,094 particles at a box size of 240 was performed, followed by a 3D classification using a mask focusing on the MacA MP domain. After classification, we noticed that one of the classes could be mixed with particles with MacB and without MacB, which gave a model at low resolution at the MacA MP domain. However, the other class showed a higher resolution feature at the MacA MP domain without the density of MacB. Further refinement was done by using the particles from this class under a C3 symmetry, and a final model was obtained around 12 Å.

### MIC assay

MIC of erythromycin was determined by a two-fold method as described previously with minor modifications^21^. Briefly, exponentially growing *E. coli* cultures (OD_600_ of 0.8) were inoculated at a density of 10^4^ cells per ml into an LB medium containing protein expression inducers and appropriate antibiotics in the presence of serial twofold increasing concentrations of erythromycin. Cell growth was determined visually after incubation at 37°C for 20 h. For the *E. coli* BW25113-Δ*acrB* cells expressing Bpa variants, cultures were inoculated at a density of 10^6^ cells per ml, as the solvent sodium hydroxide used for dissolving Bpa is toxic to bacteria.

### Formaldehyde-mediated *in vivo* chemical cross-linking

Plasmids pBAD-*macAB-his* and pRSF-*tolC* were co-transformed into BL21(DE3) competent cells to overexpress MacA, MacB, and TolC. After induction, *E. coli* cells were mixed with formaldehyde (at a final concentration of 0.1% (v/v)) at room temperature for cross-linking (0 min, 5 min, 10 min, 15 min) before quenching with glycine (at a final concentration of 0.5 M). Cells were harvested and lysed overnight with SDS sample loading buffer before being subjected to SDS-PAGE and immunoblot analysis using anti-His tag monoclonal antibody (1:1000; Cat# HRP-66005, Proteintech).

### *In vivo* site-specific photo-crosslinking mediated by photo-reactive amino acid Bpa

For Bpa-mediated *in vivo* photo-crosslinking experiments, plasmids for expressing Bpa variants of MacA and MacB were generated from the template plasmids pBAD-*macAB*-his, using the QuickChange site-directed mutagenesis kit (Transgene, Beijing, China). The pSup-BpaRS6TRN plasmid (chloramphenicol resistant), which expresses the orthogonal aminoacyl-tRNA synthetase/tRNA pair for the incorporation of Bpa into the protein of interest, was preserved in our lab and was co-transformed with pBAD-*macAB*-his and pRSF-*tolC* into *E. coli* BL21(DE3)-Δ*acrAB-tolC* cells. Cells were cultured in the presence of 100 *µ*g/ml ampicillin, 50 *µ*g/ml kanamycin, 50 *µ*g/ml chloramphenicol, 1 mM Bpa (Macklin, Shanghai, China), and 0.02% L-arabinose and 0.1 mM IPTG to induce protein expression.

*E. coli* cells were transferred to a 6-well plate (on ice) followed by UV irradiation at 365 nm for appropriate time using an ultraviolet cross-linker (Scientz, Ningbo, China). The cells were then lysed with SDS sample loading buffer before being analyzed by SDS-PAGE and immunoblotting with anti-His tag monoclonal antibody (1:1000; Cat# HRP-66005, Proteintech).

### Semi-quantification of relative MacA-MacB cross-linked product

The relative level of photo-crosslinked MacA-MacB was calculated as the percentage of MacA-MacB to the sum of MacA-MacB and MacB/MacA monomer based on the immunoblotting results. GraphPad Prism 9.0 was used for data analysis and figure generation. Western blotting data were analyzed with T-tests. Differences were considered significant when *P* < 0.05. Data are shown as the mean values ± SD.

## Acknowledgments

This work was supported by R01GM143380, R01HL162842, and Welch Foundation Q-2173-20230405 funds to Z.W.; the National Natural Science Foundation of China (No. 82072312), and the Natural Science Foundation of Jiangsu Province (No. BK20211053) to X.S.; Postgraduate Research & Practice Innovation Program of Jiangsu Province (No. KYCX22_2925 and No. KYCX23_2958); and NIH R01GM080139 to S.J.L. Cryo-EM data was collected at the Baylor College of Medicine Cryo-EM ATC and UTHealth cryo-EM Core, which includes equipment purchased under the support of CPRIT Core Facility Award RP190602.

## Author contributions

X.S., Z.W., and T.H. designed experiments. Z.Y. and X.S. performed sample freezing and cryo-ET imaging. T.H. and Z.Y. did cryo-ET data processing. W.Z., Y.W., and Q.R. performed biochemistry experiments. T.H. and X.S. wrote the initial manuscript. Z.W., J.C., and Z.Y. revised the manuscript.

## Supplementary Figures

**Supplementary Figure 1.**
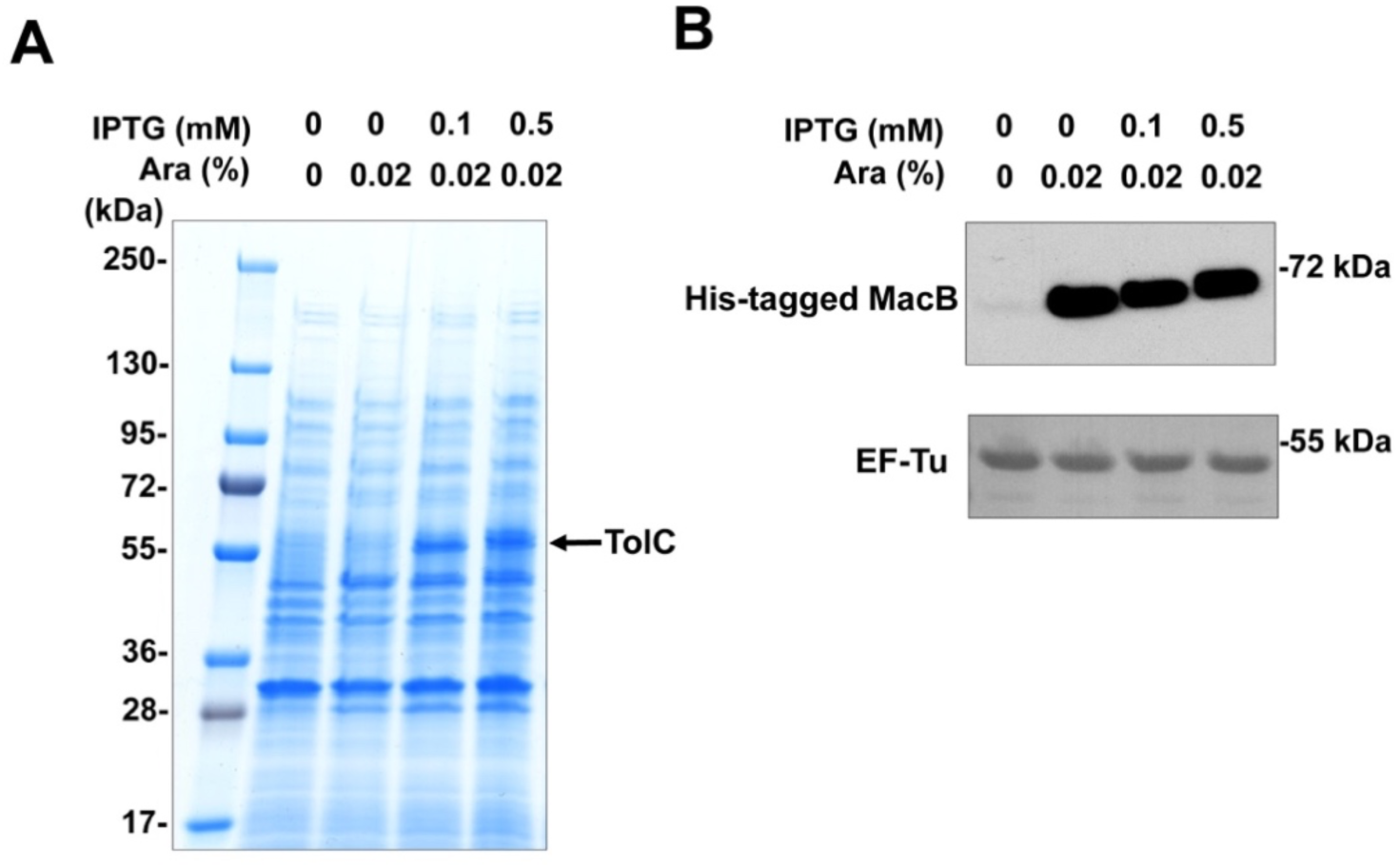
Determination of the overexpression of MacA, MacB, and TolC in BL21(DE3) cells. Plasmids pBAD-*macAB-his* and pRSF-*tolC* were co-transformed into BL21 (DE3) competent cells. Protein was induced by IPTG and L-arabinose at 20°C overnight. A, Coomassie Brilliant Blue staining results of TolC overexpression. B, immunoblotting results of MacB overexpression by using anti-his antibodies. The expression of EF-Tu was detected by anti-EF-Tu antibodies as the internal control. Ara, arabinose.

**Supplementary Figure 2.**
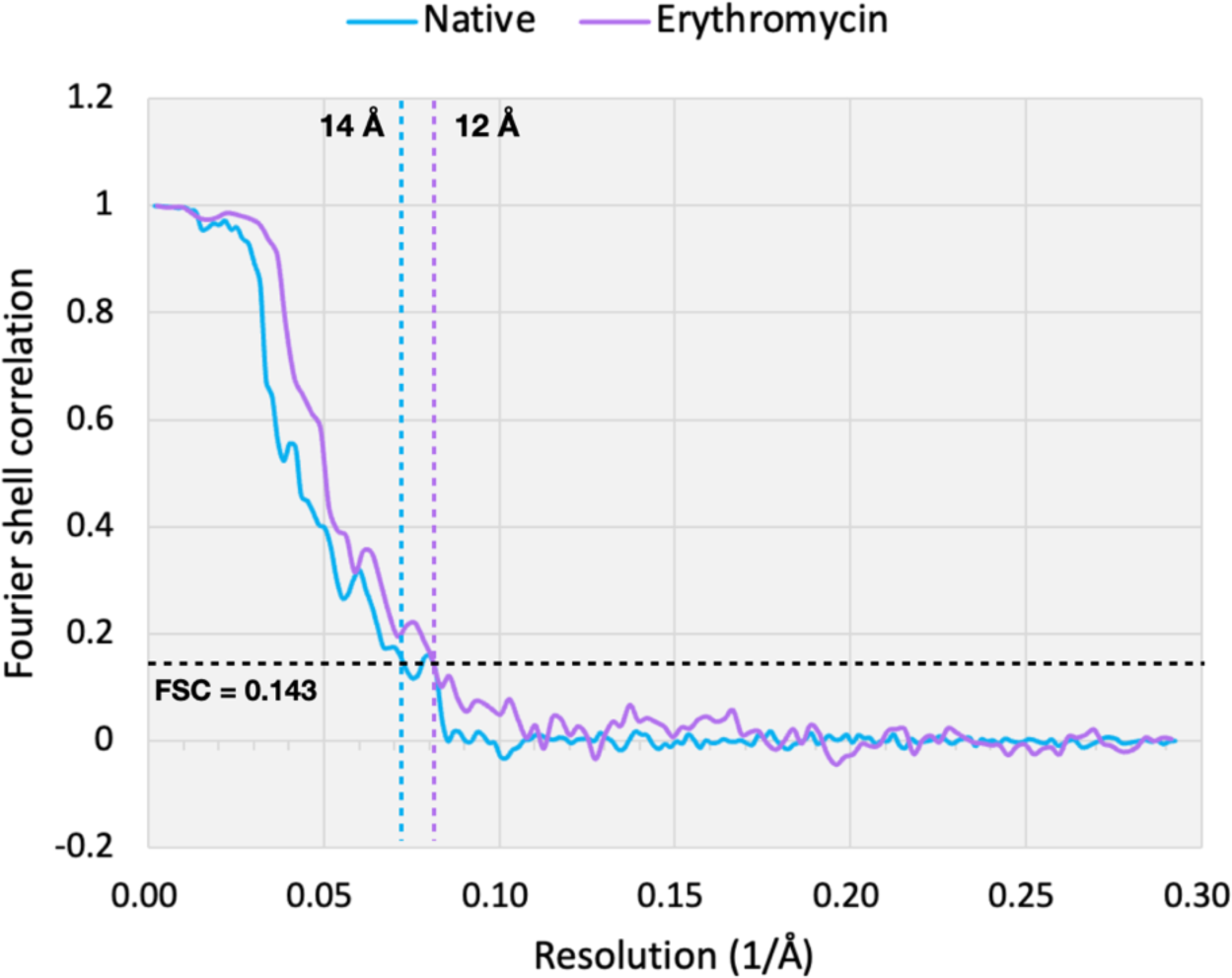
The FSC curves show the resolution of the structures achieved in this work (FSC threshold equals 0.143).

**Supplementary Figure 3.**
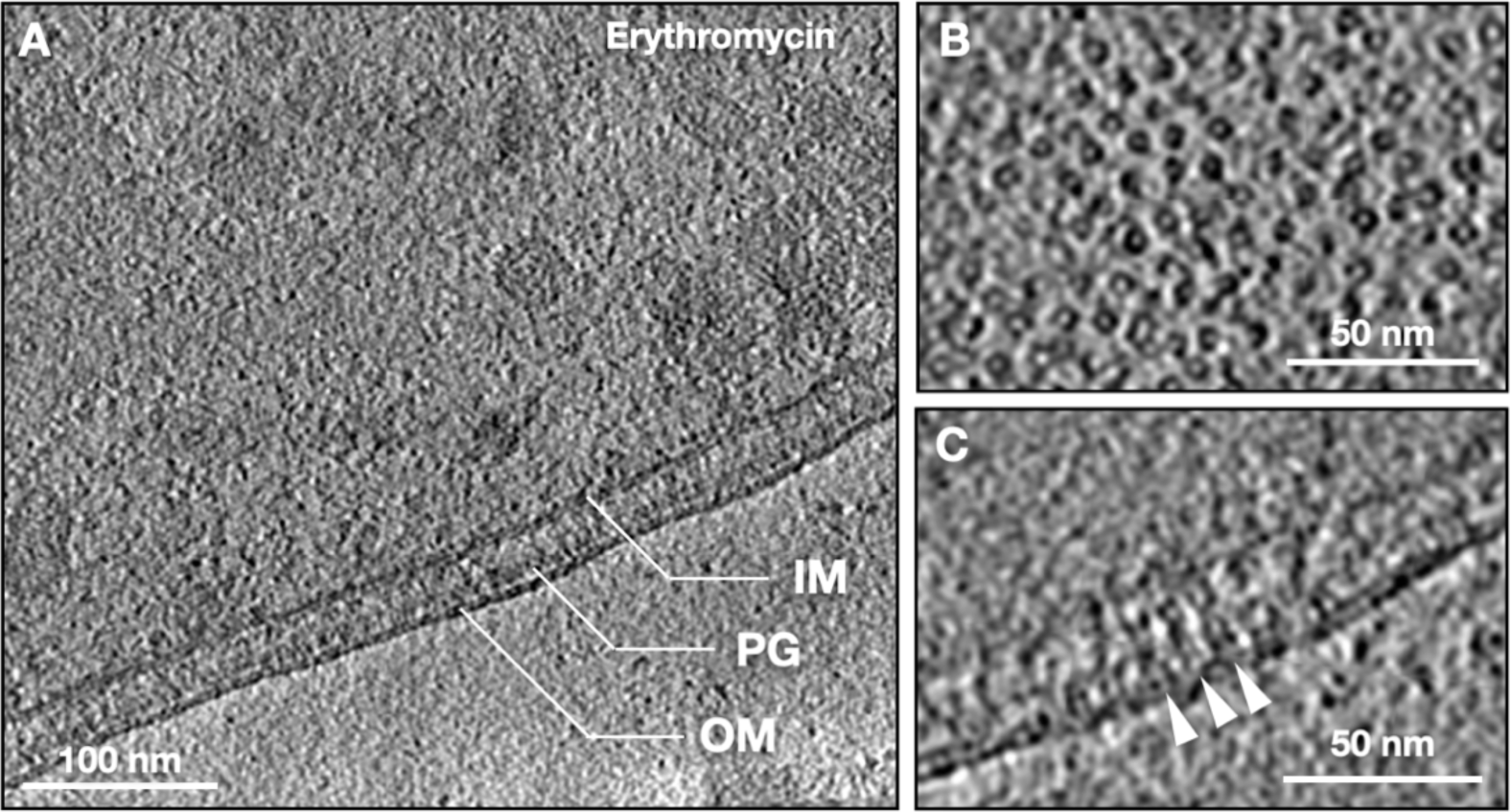
The MacAB-TolC pumps visualized in *E. coli* cells with erythromycin treatment. A, a single Z slice from a tomogram of an *E. coli* cell expressing MacA, MacB, and TolC and added with erythromycin. B, zoom-in view of the MacAB-TolC top view particles. C, zoom-in view of the MacAB-TolC side view particles (indicated by white arrowheads). IM, inner membrane; OM, outer membrane; PG, peptidoglycan.

**Supplementary Figure 4.**
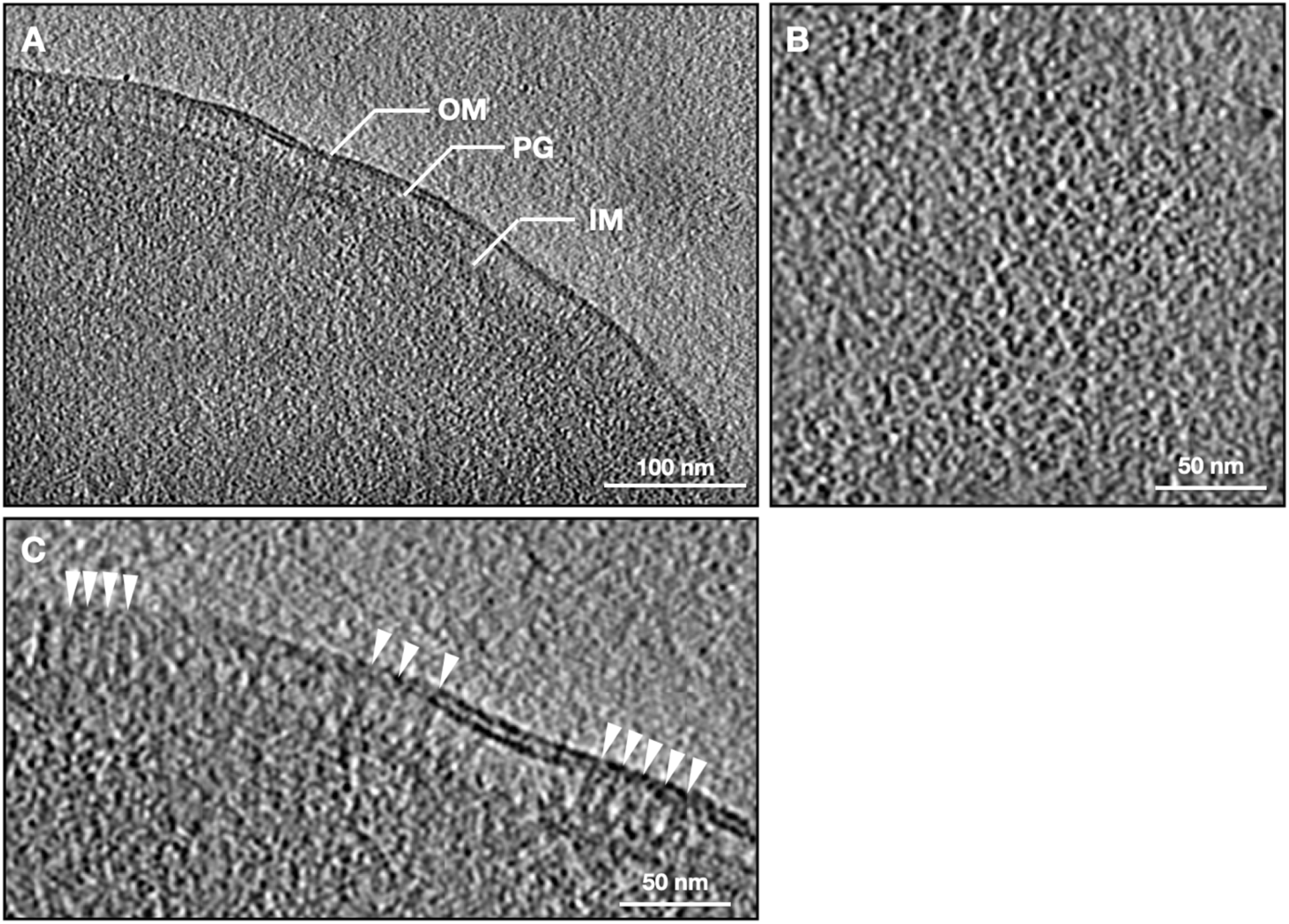
The MacA-TolC subcomplex visualized in *E. coli* cells. A, a single Z slice from a tomogram of an *E. coli* cell expressing MacA and TolC. B, zoom-in view of the MacA-TolC top view particles. C, zoom-in view of the MacA-TolC side view particles (indicated by white arrowheads). IM, inner membrane; OM, outer membrane; PG, peptidoglycan.

**Supplementary Figure 5.**
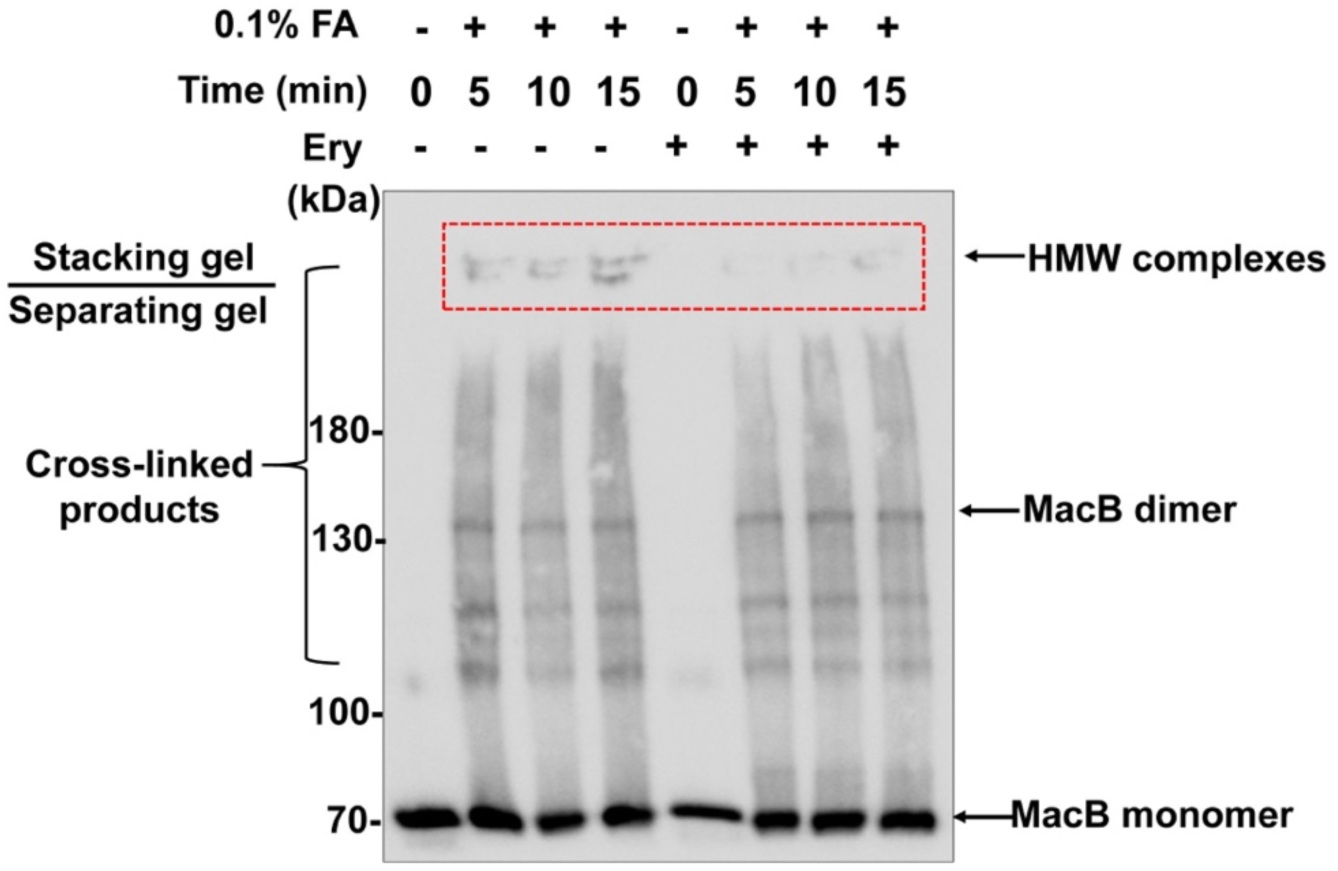
Immunoblot results with formaldehyde-mediated *in vivo* cross-linked products of MacB upon erythromycin treatment. *E. coli* cells overexpressing the MacAB-TolC efflux pump were treated with/without erythromycin (400 *µ*g/ml) at room temperature for half an hour and then mixed with formaldehyde (0.1%) for cross-linking before quenching with glycine (0.5 M). Cells were lysed before being subjected to SDS-PAGE and immunoblot analysis. FA, formaldehyde; HMW, high molecular weight; Ery, erythromycin.

**Supplementary Figure 6.**
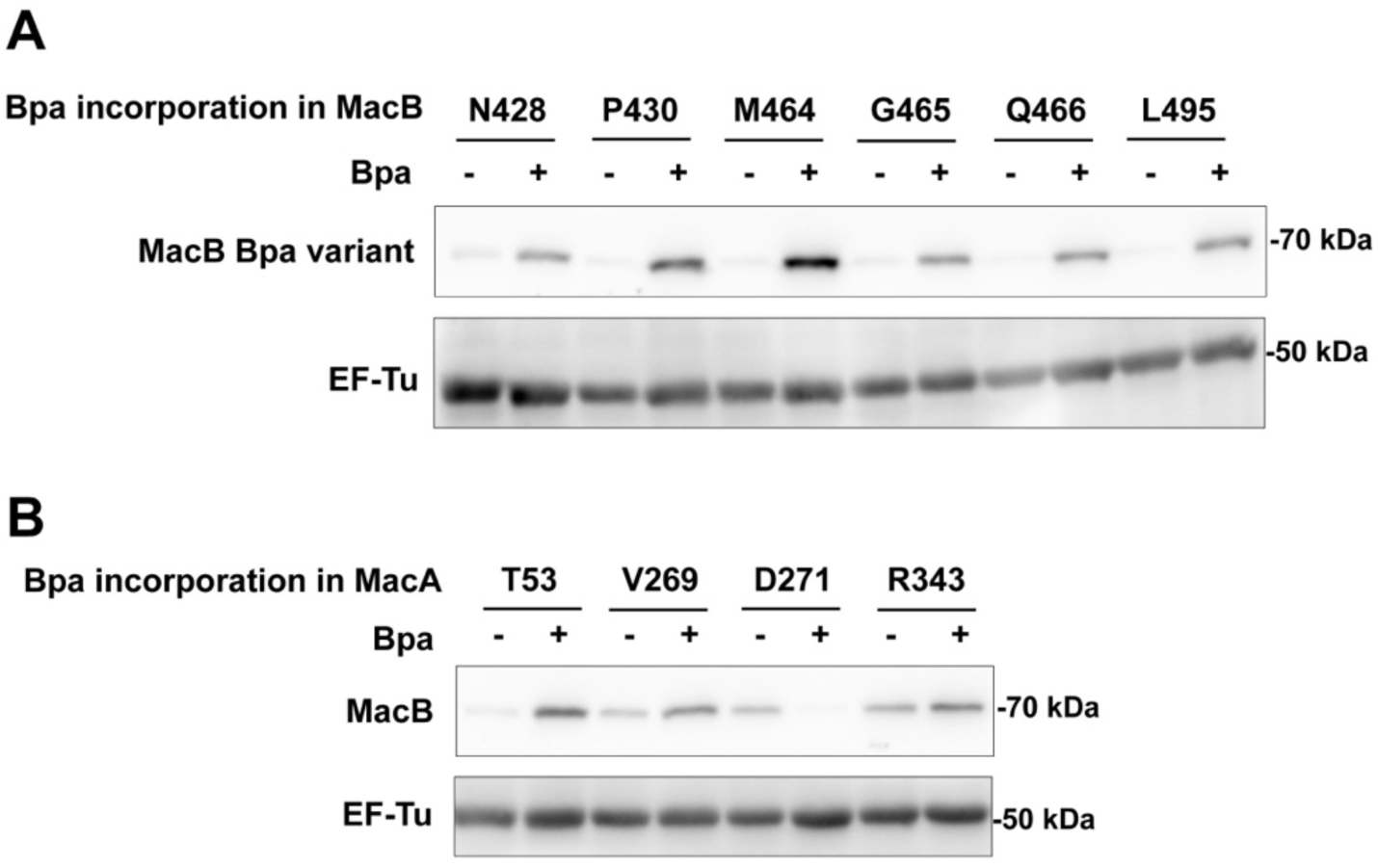
Determination of the expression of MacB Bpa variant/MacB. Protein was induced by 0.02% L-arabinose in the absence/presence of Bpa. The expression of MacB Bpa variant/MacB was detected by anti-His tag monoclonal antibodies. The expression of EF-Tu was detected by anti-EF-Tu monoclonal antibodies as the internal control. A, immunoblotting results of six MacB Bpa variants. B, immunoblotting results of the expression of MacB after the Bpa being incorporated in MacA at four different selected individual positions.

**Supplementary Figure 7.**
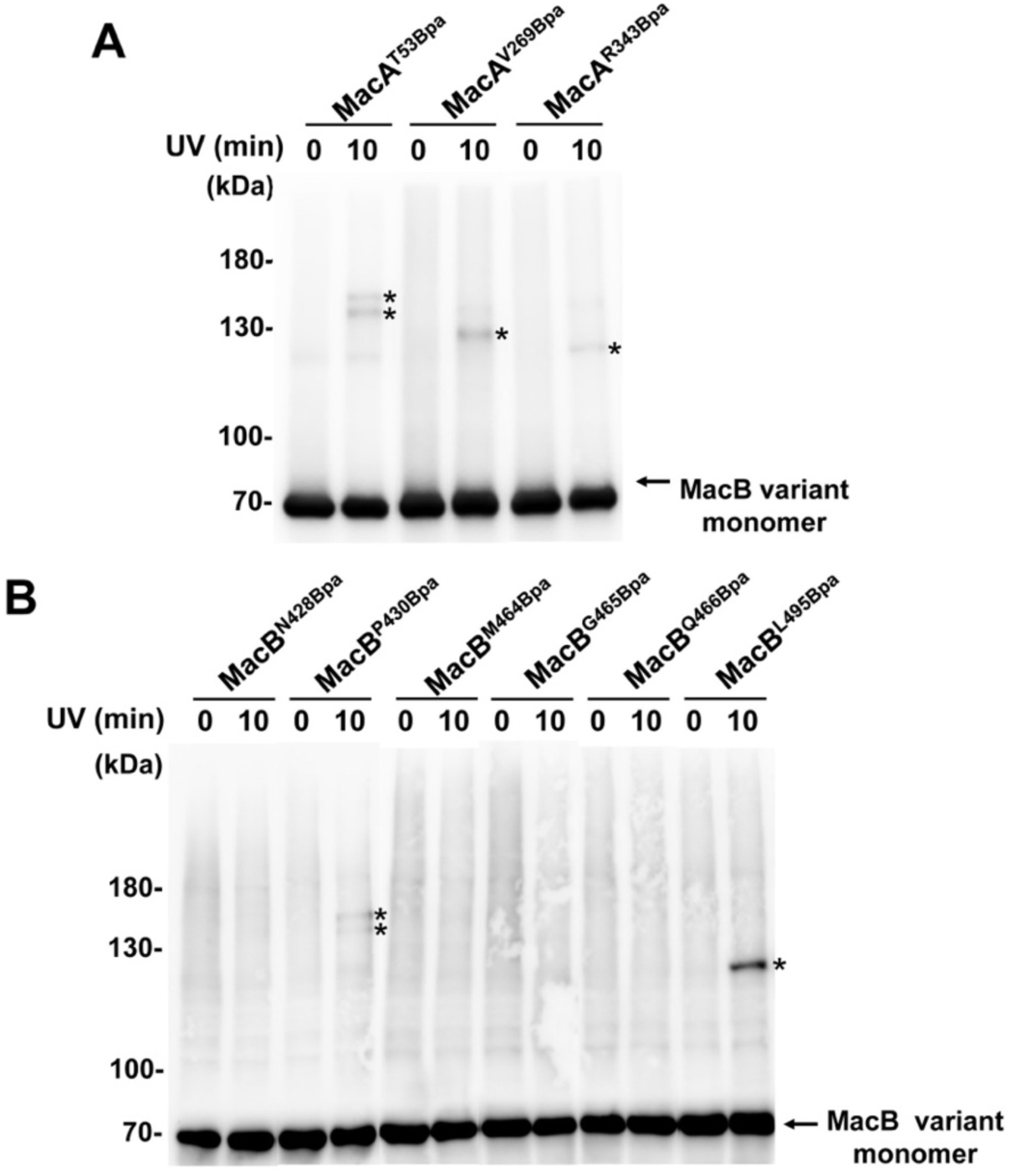
Immunoblotting results of the *in vivo* photo-crosslinked products of all the MacA/MacB Bpa variants. A, the photo-crosslinking results of the MacA Bpa variants. B, the photo-crosslinking results of the MacB Bpa variants. The bands indicated by the asterisk represent the cross-linked products of MacA Bpa variants with MacB or MacA with MacB Bpa variants.

## Supplementary Tables

**Supplementary Table 1.** Erythromycin susceptibility of *E. coli* BL21(DE3)-Δ*acrAB-tolC* cells overexpressing MacA, MacB, and TolC.

| Protein expressed | MIC (μg/ml) |
| --- | --- |
| — | 1 |
| MacA and MacB | 1 |
| MacA, MacB, and TolC | 4 |

Note: Plasmids pBAD-*macAB-his* was transformed into BL21(DE3)- $\Delta$ *acrAB-tolC* competent cells to overexpress MacA and MacB. Plasmids pBAD-*macAB-his* and pRSF-*tolC* were co-transformed into BL21(DE3)- $\Delta$ *acrAB-tolC* competent cells to overexpress MacA, MacB, and TolC. Protein was induced by 0.5 mM IPTG (for TolC) and 0.2% L-arabinose (for MacA and MacB).

**Supplementary Table 2.** Erythromycin susceptibility of BW25113-Δ*acrB* cells overexpressing MacA/MacB Bpa variants.

| Protein expressed | MIC (μg/ml) |
| --- | --- |
| — | 2 |
| MacA, MacB | 4 |
| MacA<sup>T53Bpa</sup>, MacB | 4-8 |
| MacA<sup>V269Bpa</sup>, MacB | 4-8 |
| MacA<sup>R343Bpa</sup>, MacB | 4 |
| MacA, MacB<sup>P430Bpa</sup> | 4-8 |
| MacA, MacB<sup>L495Bpa</sup> | 4 |

**Supplementary Table 3.** Statistics of *in situ* structures achieved in this study.

| Structure | MacAB-TolC<br>(native) | MacAB-TolC<br>(Erythromycin) | MacA-TolC |
| --- | --- | --- | --- |
| Overexpressed protein(s) | MacA, MacB, TolC | MacA, MacB, TolC | MacA, TolC |
| Treatment | N/A | Erythromycin | N/A |
| **Microscope** | Titan Krios | Titan Krios | Titan Krios |
| **Tilt scheme** | Bidirectional | Bidirectional | Bidirectional |
| **Angstrom per pixel** | 1.76 | 1.76 | 1.76 |
| **Tilt series number** | 70 | 68 | 67 |
| **Particle number** | 5,183 | 1,438 | N/A |
| **Resolution (Å)** | 14 | 12 | N/A |

